# Inhibition of Histone deacetylase 1 (HDAC1) and HDAC2 enhances CRISPR/Cas9 genome editing

**DOI:** 10.1101/670554

**Authors:** Bin Liu, Siwei Chen, Anouk La Rose, Deng Chen, Fangyuan Cao, Dominik Kiemel, Manon Aïssi, FJ Dekker, HJ Haisma

**Author notes:** Bin Liu and Siwei Chen equally contributed to this manuscript. Corresponding author Hidde J. Haisma, Department of Chemical and Pharmaceutical Biology, Groningen Research Institute of Pharmacy, University of Groningen, 9713 AV Groningen, The Netherlands.

## Abstract

Despite the rapid development of CRISPR/Cas9-mediated gene editing technology, the gene editing potential of CRISPR/Cas9 is hampered by low efficiency, especially for clinical applications. One of the major challenges is that chromatin compaction inevitably limits the Cas9 protein access to the target DNA. However, chromatin compaction is precisely regulated by histone acetylation and deacetylation. To overcome these challenges, we have comprehensively assessed the impacts of histone modifiers such as HDAC (1-9) inhibitors and HAT (p300/CBP, Tip60 and MOZ) inhibitors, on CRISPR/Cas9 mediated gene editing efficiency. Our findings demonstrate that attenuation of HDAC1, HDAC2 activity, but not other HDACs, enhances CRISPR/Cas9-mediated gene knockout frequencies by NHEJ as well as gene knock-in by HDR. Conversely, inhibition of HDAC3 decreases gene editing frequencies. Furthermore, our study showed that attenuation of HDAC1, HDAC2 activity leads to an open chromatin state, facilitates Cas9 access and binding to the targeted DNA and increases the gene editing frequencies. This approach can be applied to other nucleases, such as ZFN and TALEN.

## Introduction

CRISPR/Cas9 is derived from the bacterial immune system where it disrupts foreign genetic elements invaded from plasmids and phages, which are eventually naked DNA. Nowadays, it is widely used in genome editing for eukaryotes, including humans. However, the eukaryotic chromosomes are more complex than their prokaryotic counterparts. In eukaryotes, DNA is packed into chromosomes in the cell nucleus in a highly compact and organized manner named chromatin. The chromatin is made up of repeating units called nucleosomes. The nucleosome consists of 147 base pairs wrapped around histone protein octamers H2A, H2B, H3, and H4. Thus, the gene editing process of CRISPR/Cas9 in eukaryotes is very different as compared to the prokaryotic process.

CRISPR/Cas9 system is revolutionizing the field of biochemical research, but a higher efficiency is anticipated for clinical practice. The efficiency of genome editing by CRISPR/Cas9 varies from 2% to approximately 25% depending on the cell type^1^, which is not yet up to the requirements for clinical use, such as cancer gene therapy^2^. Most approaches for optimizing CRISPR based techniques are mainly focused on optimizing the structure of gRNAs^3–5^, creating mutant Cas9^6^ and finding new versions of CRISPR/Cas system from prokaryotes^7,8^, etc. Although these approaches are essential, the underlying genomic context, particularly the chromatin state of the target locus, significantly influences the cleavage efficiency^9,10^. Recent studies showed that the targeting efficiency of CRISPR/Cas9 varied widely in different target loci of the chromosome^10,11^. The euchromatic target sites show higher frequencies of DSB (double-strand break) introduced by TALENs and CRISPR/Cas9 as compared to those of the heterochromatic sites. Notably, a recent study showed that the spontaneous breathing of nucleosomal DNA and chromatin remodelling facilitates Cas9 to effectively act on chromatin^12^. Thus, the chromatin conformations can significantly impact gene editing efficiency of nucleases.

Undoubtedly, there is a considerable number of target sites inevitably located in heterochromatin, which has a strong effect on the accessibility of DNA to Cas9^13^. Furthermore, albeit many genes are located in a relatively euchromatic position, the gene editing efficiency may also be enhanced through maintaining the open state of those euchromatic regions. But the approaches on how to manipulate the chromatin state and efficiently target those genes in heterochromatin sites are lacking. The open or closed state of chromatin structure is mainly controlled by the balance of histone acetylation and deacetylation which is strictly regulated by two groups of enzymes called histone acetyltransferase (HAT) and histone deacetylase (HDAC). Briefly, histone acetylation leads to a loose or uncoiling of the chromatin structure (euchromatin). Conversely, histone deacetylation leads to a condensed or closed chromatin structure (heterochromatin). The euchromatin gives the transcriptional machinery access to the transcriptionally active DNA, which also provides a great opportunity for CRISPR/Cas9 attacking and cutting the DNA, particularly for the targets located in condensed heterochromatin regions. More importantly, the chromatin state regulated by HAT and HDAC may also have the potential to influence the gene knock-in mediated by HDR, which has extremely low efficiency and needs to be improved^1,14^. In addition, previous studies showed that the dCas9 fused to core p300 or HDAC3 robustly influences epigenome editing^15,16^, but the effects of these HATs or HDACs on genome editing of CRISPR/Cas9 have yet to be characterized.

Given the development of histone modifiers such as HAT, HDAC inhibitors and other biotechnology approaches^17^, it is possible and rational to explore whether the gene editing efficiency can be improved by altering the chromatin state through modulation of the HDAC and HAT activity. We hypothesized that the regulation of chromatin compaction by inhibiting HAT and/or HDAC activity can modulate CRISPR/Cas9 based gene editing. Our findings show that inhibition of HDAC1, HDAC2, rather than other HDACs, can enhance both gene knockout and gene knock-in. We also show that inhibition of HDAC3 could decrease the efficiency of CRISPR/Cas9 mediated gene editing. Furthermore, we provide a practical and clinically applicable approach for precise control of CRISPR/Cas9 mediated gene editing by modulation of HDAC and HAT activity in host cells.

## Results

### 1. The landscape of HDAC/HAT inhibition on CRISPR/Cas9-mediated gene knockout

To investigate the effect of different HDACs/HATs on CRISPR/Cas9-mediated gene knockout, we tested a panel of HDAC/HAT inhibitors (**Table 1**) with a CRISPR/Cas9-mediated EGFP knockout reporter system^18^ (**Figure 1 A**). To preclude the interference of drug toxicity, we performed a cell viability assay with the treatment of HDAC/HAT inhibitors in H27 and HT29 cells (**Figure S1**). At least 90% of cell viability compared to the control group was considered as acceptable. According to the cell viability data (**Figure S1**), the dose of each inhibitor without affecting cell viability are Panobinostat (0.1 µM), Entinostat (5 µM), RGFP966 (1 µM), TMP195 (1 µM), PCI34051 (1 µM), Tubastatin A (1 µM), C646 (1 µM) and MG149 (1 µM) (**Figure S1 A**).

**Table 1.**
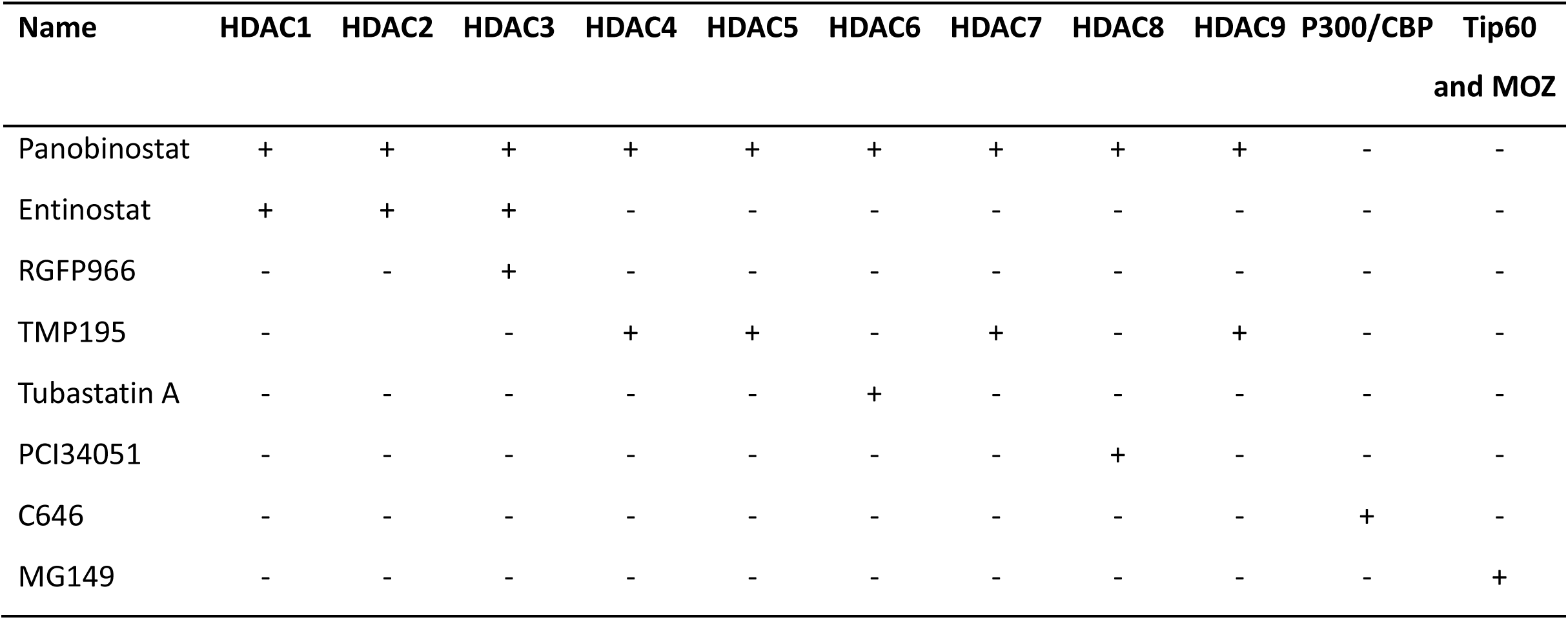
List of HDAC/HAT inhibitors used in this study.

**Figure 1.**
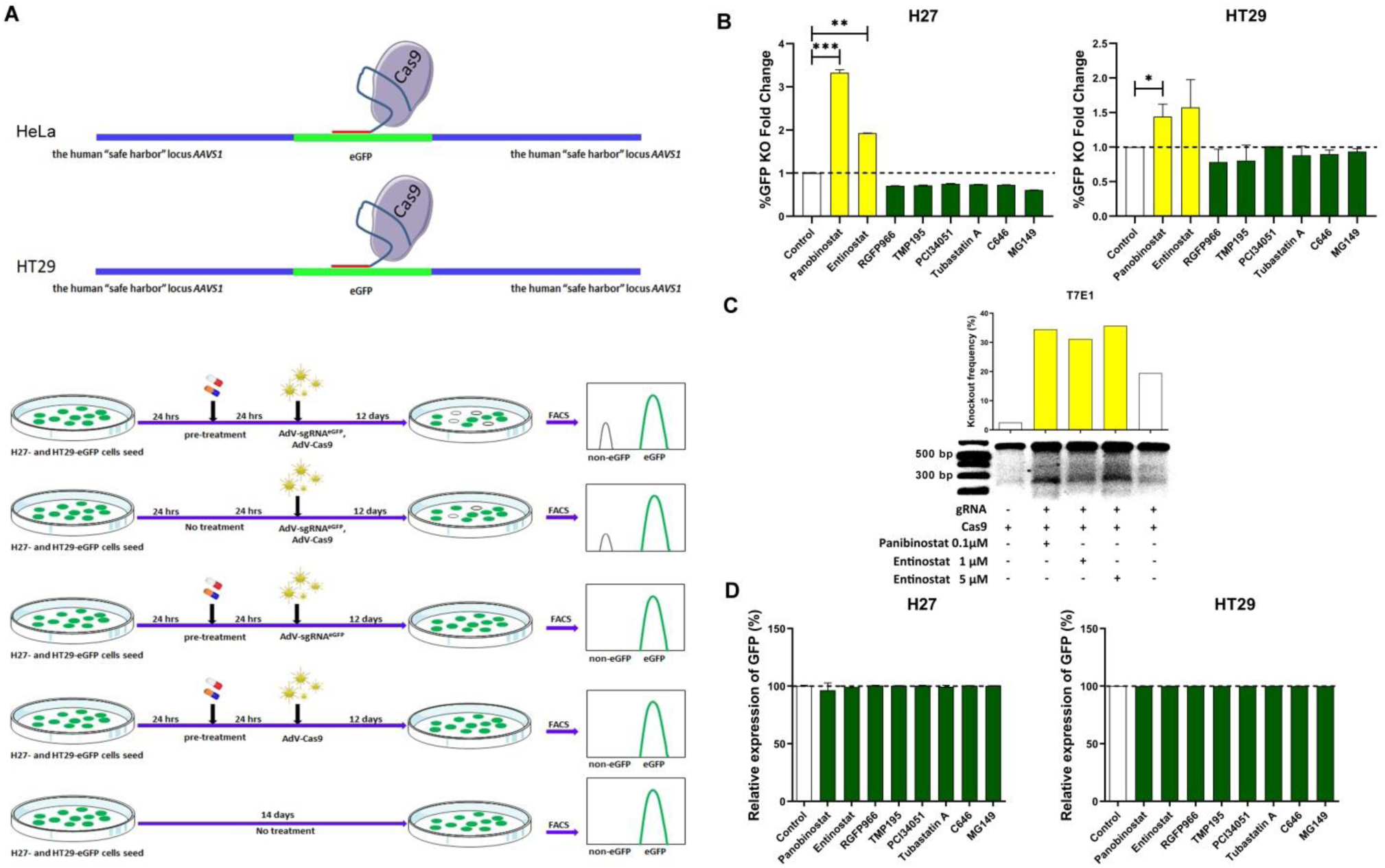
Screening HDAC inhibitors for improving CRISPR/Cas9 mediated gene editing. The work flow of HDAC/HAT inhibitors screening and their effects on CRISPR/Cas9 mediated gene knockout. **A.** Schematics of HDAC inhibitors screening procedures for CRISPR/Cas9 mediated gene knockout. CRISPR/Cas9 specifically cuts the EGFP sequence in HeLa (H27) and HT29 cells resulting in EGFP signal loss. Cells were pre-treated with different inhibitors and incubated with AdV-Cas9 and AdV-gRNA, subsequently subcultured for 12 days to remove EGFP protein from cells with disrupted EGFP ORFs. **B.** The effect of different HAT/HDAC inhibitors on CRISPR/Cas9 mediated gene knockout in H27 and HT29. The frequencies of gene knockout were quantified by flow cytometry. **C.** T7 endonuclease I (T7EI) genotyping assays for detection of indels induced by CRISPR/Cas9 and quantified by ImageJ. **D.** Endogenous EGFP expression determined by flow cytometry with the treatment of different HDAC/HAT inhibitors. Data in bar graphs are represented as mean ± SD (n≥3), two-tailed unpaired student’s t-test: *p-values < 0.05; **p-values < 0.01; ***p-values < 0.001.

A co-transduction grid experiment was performed to assess which dosage of virally delivered gRNA and Cas9 results in high genomic EGFP gene knockout levels without reducing cell viability. We observed that a dosage of 30 TU/cell resulted in relatively high amounts of EGFP gene knockout as well as no reduction of cell viability (**Figure S2**). Therefore, we transduced 1:1 mixtures of and gRNA and Cas9 viral particles at 30 TU/cell, respectively, to induce DSBs at EGFP sequences. The EGFP gene editing frequencies were determined by direct fluorescence microscopy and flow cytometry at 14 days post-transduction. Our results showed that Entinostat (HDAC1/2/3 inhibitor) and Panobinostat (pan-HDAC-inhibitor) significantly increased gene knockout frequencies in both H27 and HT29 cells by 1.5∼3.4 folds (**Figure 1 B**). Of note, RGFP966 (HDAC3 inhibitor) slightly decreased gene knockout frequencies as well as other HDAC inhibitors (HDAC4-9) (**Figure 1 B**). Additionally, HAT inhibitors C646 (p300/CBP inhibitor) and MG149 (Tip60 and MOZ inhibitor) decreased the EGFP gene knockout frequencies (**Figure 1 B**). The gene knockout frequencies enhanced by Entinostat and Panobinostat were further determined by T7 endonuclease I (T7E1) assay (**Figure 1C**). As controls, we did not observe gene knockout in the groups with a single treatment (gRNA only, Cas9 only, inhibitors only) and non-treatment groups. To assess the possibility of the impacts of HDAC inhibitors on endogenous EGFP expression in H27 and HT29, we tested the EGFP expression level using flow cytometry and there were no changes of endogenous EGFP expression, indicating no affection on endogenous EGFP expression upon these HAT/HDAC inhibitors treatments (**Figure 1D**).

### 2. Panobinostat and Entinostat enhance CRISPR/Cas9-mediated gene knockout in a dose-dependent manner

Next, we asked whether the enhancement of gene knockout by Panobinostat and Entinostat is dose dependent. We tested the indicated amounts of Panobinostat and Entinostat (**Figure 2**). Our results showed a positive correlation between the EGFP gene knockout frequencies and the dose of Panobinostat and Entinostat (**Figure 2**). These results clearly indicate that Panobinostat and Entinostat enhance CRISPR/Cas9-mediated gene knockout in a dose-dependent manner.

**Figure 2.**
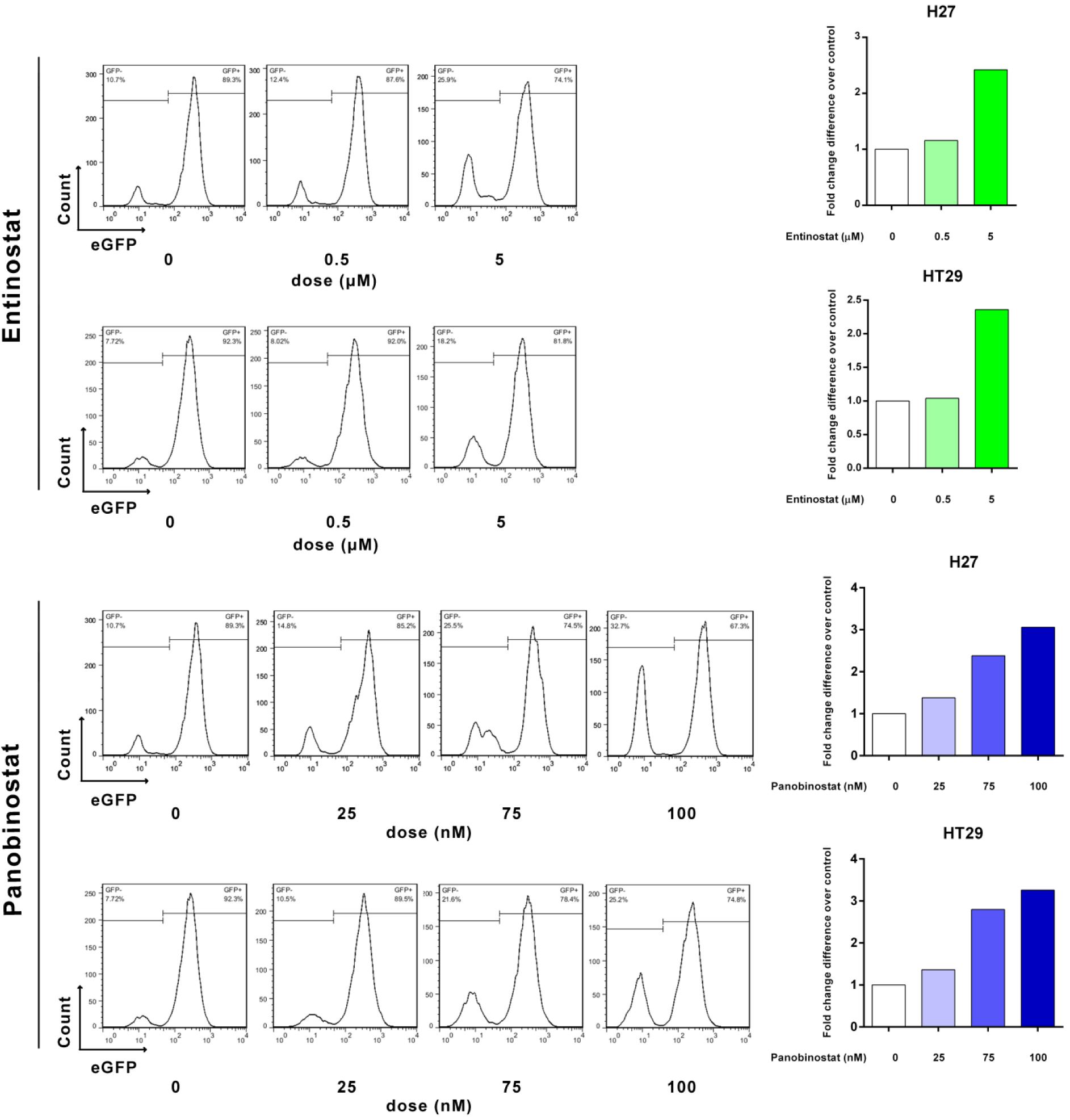
Dose dependent effects of entinostat and panobinostat on CRISPR/Cas9 mediated gene knockout. The dosage of entinostat are 0, 0.5 and 5 µM and panobinostat are 0, 25, 75, 100 nM. Dose-dependent response to Entinostat and Panobinosta was assessed by flow cytometry. Data were derived from one of three independent experiments. The quantification bar graph (right) generated according to flow cytometry data (left).

### 3. Downregulation of HDAC1 or HDAC2 increases gene knockout efficiency

Both Panobinostat (a pan-HDAC inhibitor) and Entinostat (an HDAC1/2/3 selective inhibitor) enhanced gene knockout, but RGFP966 (an HDAC3 selective inhibitor) decreased gene knockout. Moreover, we did not observe the enhancement of gene knockout by other HDAC selective inhibitors. Therefore, we speculate that HDAC1/2/3 may play an important role in the gene editing process of CRISPR/Cas9. To precisely distinguish the role of the three HDACs in gene editing processes, we used siRNAs to specifically knockdown HDAC1, HDAC2 or HDAC3 to explore whether gene knockout efficiency can be enhanced or decreased (**Figure 3A**). In agreement with the inhibitors results above, we showed that knockdown of HDAC1 or HDAC2 results in a significant increase by 16%∼28% and 22%∼26% (p<0.01) of EGFP gene knockout, respectively (**Figure 3B**). Moreover, our results showed that downregulation of HDAC3 decreased the EGFP gene knockout efficiency by 21% and 39% (p<0.001) in H27 and HT29 cell lines, respectively (**Figure 3B**). Thus, downregulation of HDAC1 or HDAC2 increases CRISPR/Cas9-mediated gene knockout, but downregulation of HDAC3 decreases it.

**Figure 3.**
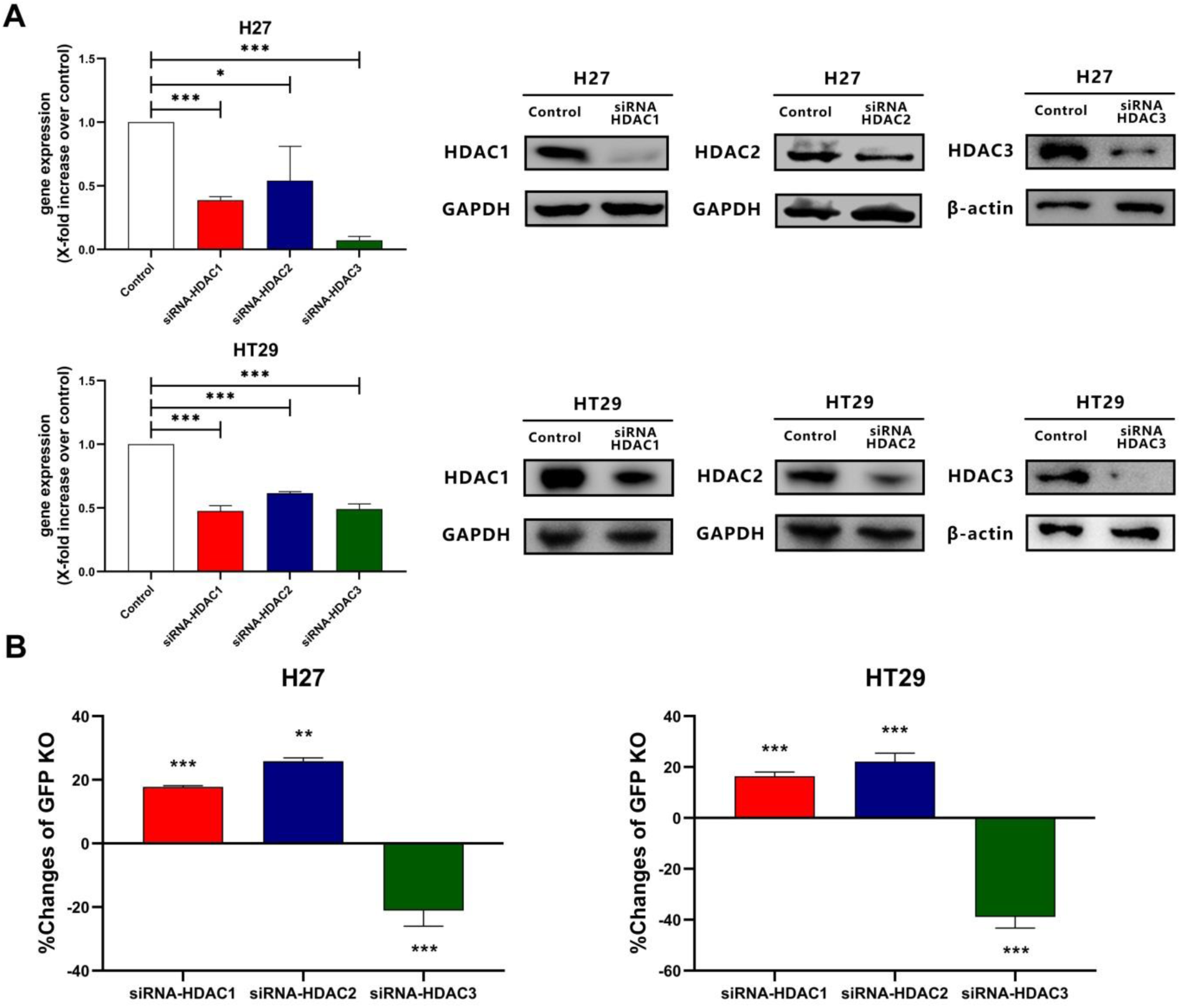
Assessment of gene knockout frequencies by knockdown of HDAC1, HDAC2 and HDAC3. **A.** HDAC1, HDAC2 and HDAC3 mRNA and protein levels were determined by RT-qPCR and western blot after transfection with scrambled siRNAs or HDAC1, HDAC2 or HDAC3 siRNAs. **B.** Changes of GFP knockout mediated by CRISPR/Cas9 upon knockdown of HDAC1, HDAC2 or HDAC3. Data in bar graphs are represented as mean ± SD (n≥3), two-tailed unpaired student’s t-test: *p-values < 0.05; **p-values < 0.01; ***p-values < 0.001.

### 4. Effect of Panobinostat and Entinostat on viral transduction, transgene transcription and cell cycle

Although we hypothesized that HDAC inhibitors may increase the CRISPR/Cas9 mediated gene editing by increasing the accessibility of the target loci, viral transduction and transgene transcription might also influence the gene knockout, particularly by affecting Cas9 protein expression levels. Therefore, additional experiments were performed to determine the influence of Panobinostat and Entinostat on viral transduction and transgene expression.

To explore the impacts of Panobinostat and Entinostat on adenovirus transduction and transgene expression, we first tested with an EGFP reporter adenovirus (AdTL). Since H27 and EGFP-HT29 cells have endogenous EGFP expression which may affect the measurement, the original HeLa and HT29 cells without EGFP expression were employed. HeLa and HT29 wild-type cells were treated with Entinostat and Panobinostat, respectively, prior and post to AdTL infection to determine adenovirus transduction and transgene expression. Subsequently, the effect of these HAT/HDAC inhibitors on the expression of GFP was evaluated by flow cytometry. The results showed that the expression of exogenous GFP by AdTL increased dramatically upon Panobinostat treatment in both pre- and post-treatment (**Figure 4A**). In contrast, in the cells treated with Entinostat there was no significant change (**Figure 4A**). These results indicate that Entinostat does not affect adenovirus transduction and transgene expression.

**Figure 4.**
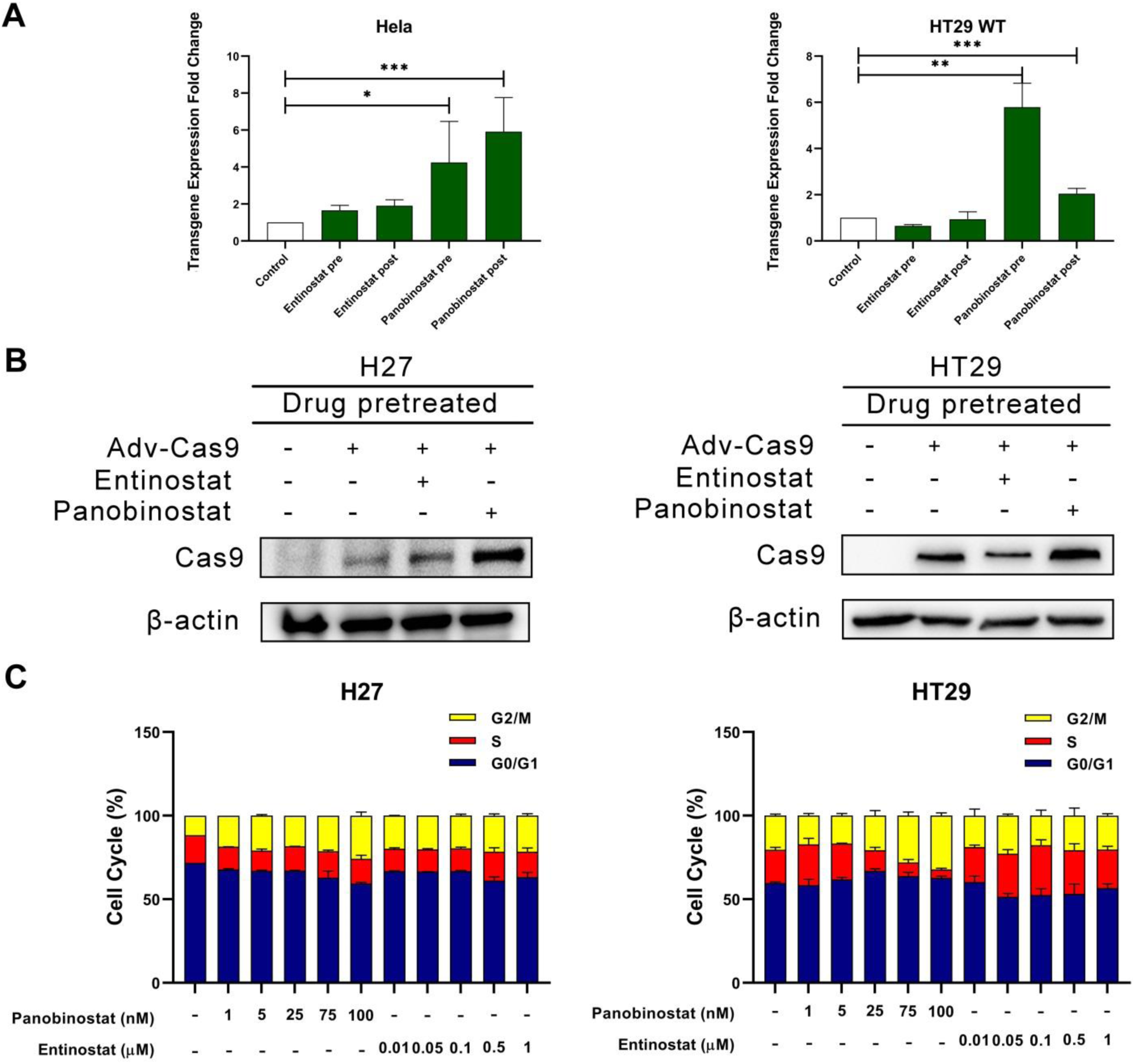
Alterations of virus transduction, transgene transcription and cell cycle with the treatment of Entinostat and Panobinostat. **A.** AdTL-EGFP transgene expression was determined by flow cytometry with the pre- and post-treatment of HDAC inhibitors. **B.** AdV-Cas9 protein expression was determined by Western blot with pre-treatment of HDAC inhibitors (Entinostat 5 µM, Panobinostat 0.1 µM). **C.** Cell cycle changes upon HDAC inhibitors treatment with different dose. Data in bar graphs are represented as mean ± SD (n≥3), two-tailed unpaired student’s t-test: *p-values < 0.05; **p-values < 0.01; ***p-values < 0.001.

To directly determine the effects of Panobinostat and Entinostat on Cas9 expression, H27 and HT29 cells were treated with Entinostat and Panobinostat, prior to AdV-Cas9 infection. Subsequently, the effects of these HAT/HDAC inhibitors on the expression of Cas9 were determined by Western blot. We observed that Panobinostat significantly increased Cas9 gene expression, but Entinostat did not show an increase of Cas9 expression (**Figure 4B**). These data indicated that the increase of gene knockout by Panobinostat may be due to Cas9 protein upregulation, but this was not the case for Entinostat. Furthermore, we checked the Cas9 expression with post-treatment. There were no obvious changes of Cas9 expression with post-treatment of Panobinostat and Entinostat (**Figure S3**). These results suggest that Panobinostat may increase Cas9 expression by enhancing viral transduction, but not transgene expression.

DSBs introduced by CRISPR/Cas9 are repaired by either non-homologous end joining (NHEJ) or homology directed repair (HDR). However, these two repairing pathways favour a specific cell cycle. NHEJ is active throughout the cell cycle, but it has the highest activity in S and G2/M stages[17], whereas HDR is most efficient in S and extremely low in G1^20^. Therefore, HDAC and HAT inhibitors might regulate the gene editing efficiency by affecting the cell cycle. To determine the cell cycle changes introduced by Entinostat and Panobinostat, we performed flow cytometry analysis of cell cycle using propidium iodide (PI) DNA staining. Panobinostat arrested cells at G2/M (0.1 µM), but we did not observe an obvious cell cycle change in the treatment of Entinostat (**Figure 4C**).

Collectively, these results show that Panobinostat and Entinostat enhance gene knockout by different mechanisms. The increase of Cas9 expression and G2/M cell cycle arrest may contribute to gene knockout enhancement induced by Panobinostat, which might be different to our initial hypothesis, whereas, Entinostat is consistent with our hypothesis.

### 5. Entinostat analogues significantly enhance the CRISPR/Cas9 gene knockout efficiency

Owing to the promising results of Entinostat for improving CRISPR/Cas9 gene editing efficiency, we tested three analogues of Entinostat to confirm our findings^21^ (**Figure 5A**). These three Entinostat analogues significantly increased the gene knockout frequencies (30% to 52%) of CRISPR/Cas9 without effects on cell viability (**Figure 5B and Figure S1 B**). In consistence with Entinostat, these analogues have no effect on endogenous EGFP expression **(Figure 5C)**, Cas9 expression (**Figure 5D and Figure S3**) and cell cycle (**Figure 5E**).

**Figure 5.**
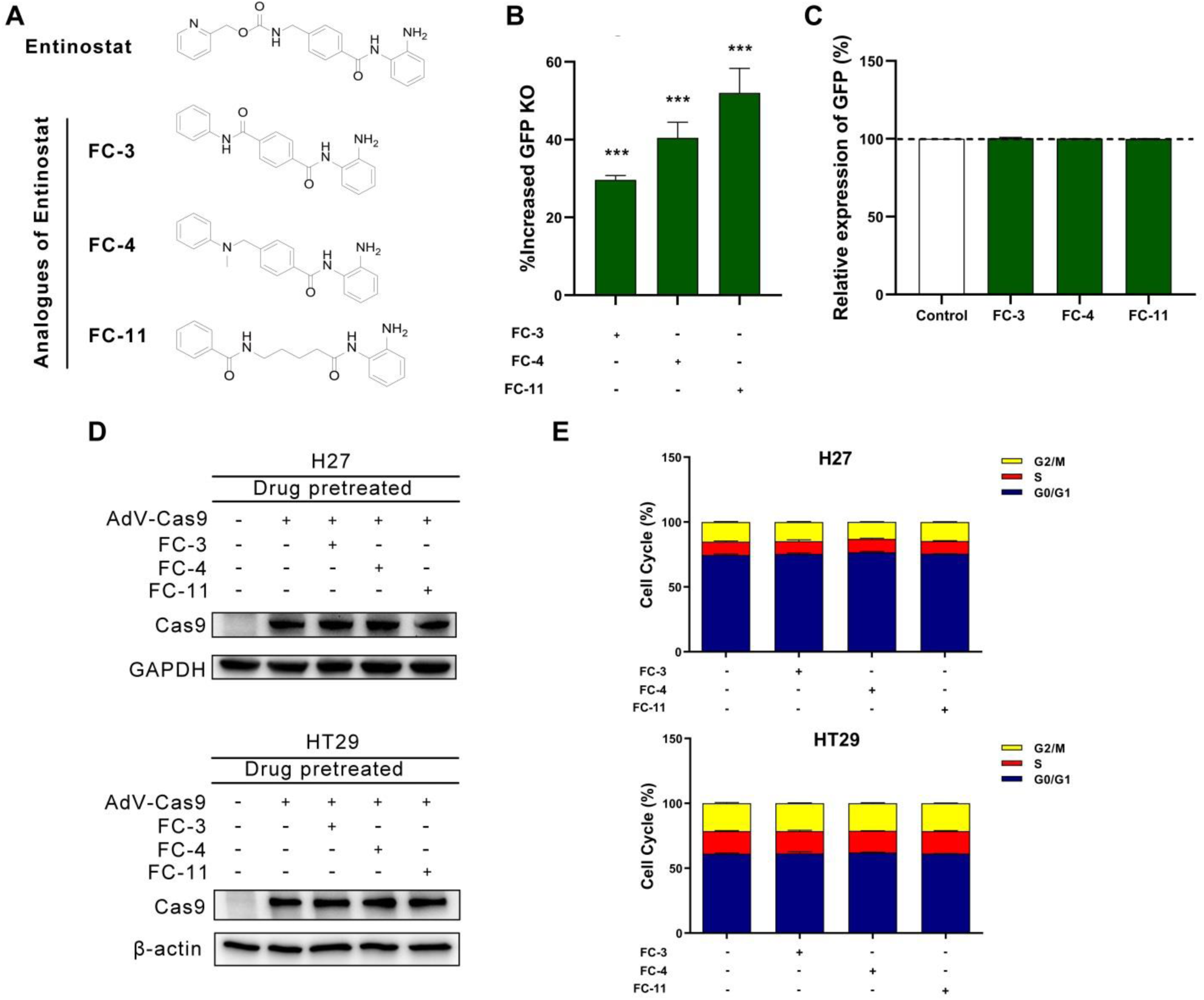
Entinostat analogues on CRISPR/Cas9 mediated gene editing. **A.** The chemical structures of Entinostat and its analogues. **B.** Gene knockout enhanced by Entinostat analogues treatment. **C.** Endogenous EGFP expression changes by Entinostat analogues treatment. **D.** Cas9 protein expression changes by Entinostat analogues treatment. **E**. Cell cycle changes by Entinostat analogues treatment. The dose of analogues used are 1 µM FC-3, 1 µM FC-4, and 10 µM FC-11. Data in bar graphs are represented as mean ± SD (n≥3), two-tailed unpaired student’s t-test: *p-values < 0.05; **p-values < 0.01; ***p-values < 0.001.

### 6. Panobinostat and Entinostat significantly enhance gene knock-in (HDR) and knockout (NHEJ)

The gene knock-in efficiency by HDR pathway is much lower than gene knockout by NHEJ. To investigate whether our inhibitors can enhance the gene knock-in efficiency, an EGFP-EBFP converting fluorescent system was employed. This system allows the simultaneous detection of NHEJ and HDR events (Figure 6A). In parallel, the chromatin compaction can be precisely regulated by doxycycline (Dox) treatment. The heterochromatin will be formed in the targeted sequences in the absence of Dox. After the gene editing process, Dox is added to determine the frequencies of NHEJ and HDR events through dual-color flow cytometry (**Figure 6A**).

**Figure 6.**
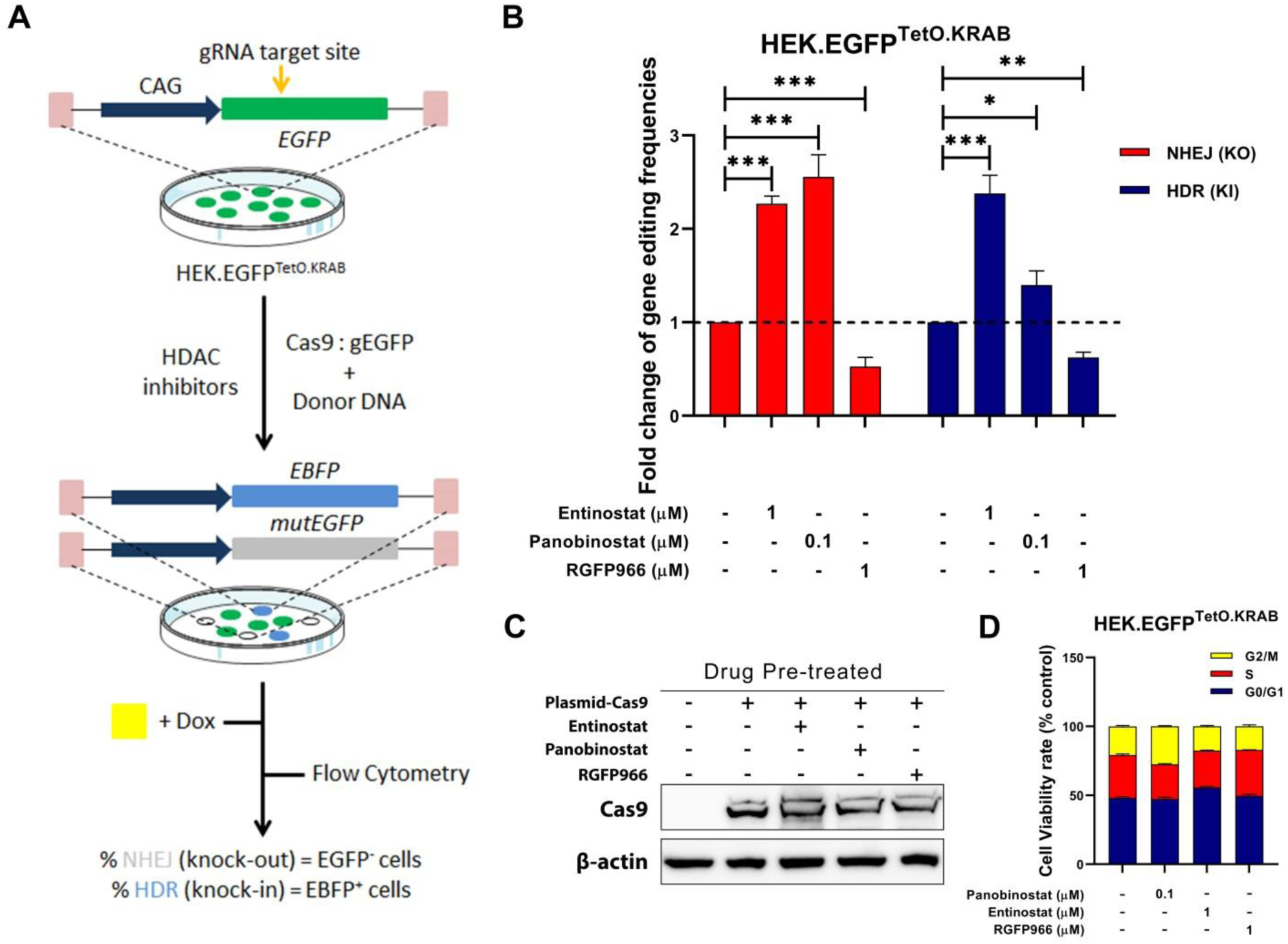
Gene-editing (NHEJ and HDR) based on converting EGFP to EBFP fluorochrome with a plasmids transient transfection system. **A.** Schematic representation of a knockout and knock-in system by converting EGFP to EBFP **B.** Alterations of gene knockout and knock-in with different HDAC inhibitors treatment **C.** Cas9 protein expression changes with different HDAC inhibitors treatment using the same dose as shown in knock-in. Data in bar graphs are represented as mean ± SD (n≥3), two-tailed unpaired student’s t-test: *p-values < 0.05; **p-values < 0.01; ***p-values < 0.001.

Entinostat increased the NHEJ and HDR rate by ∼2.3 folds, ∼2.4 folds, respectively (**Figure 6B**). Panobinostat increased the NHEJ and HDR rate by ∼2.6 folds, ∼ 1.4 folds, respectively (**Figure 6B**). However, RGFP966 decreased both knockout and knock-in. In agreement with the data from the adenovirus system, alterations of gene knockout with this plasmid transient transfection system were similar, which confirms our previous data. Meanwhile, we performed a cell viability assay to determine that the concentrations of the compounds we used have no effect on cell proliferation (**Figure S1 C**).

We further determined the effects of Panobinostat and Entinostat on Cas9 plasmid expression by Western blot. We did not observe changes of Cas9 expression levels after treatment with either Panobinostat or Entinostat (**Figure 6C**). These results suggest that the enhancement of gene knock-in/knockout by Panobinostat or Entinostat is not due to Cas9 protein changes. In addition, we also checked the Cas9 expression with Panobinostat or Entinostat post-treatment. Panobinostat slightly enhanced Cas9 protein level with post-treatment in H27, but this was not the case for Entinostat (**Figure S4)**. Furthermore, we checked the Cas9 expression with pre- and post-treatment of all Entinostat analogues **(Figure 5A)** and found that there were no increases of Cas9 expression (**Figure S4)**. Furthermore, Panobinostat and Entinostat showed no significant effect on the cell cycle of HEK.EGFP^TetO.KRAB^.

### 7. Chromatin immunoprecipitation (ChIP)-qPCR revealed Entinostat and Panobinostat introduced an open state of Chromatin in the target loci

To gain insight into the chromatin state at the CRISPR/Cas9 target region, ChIP-qPCR assays were performed in the presence or absence of Entinostat and Panobinostat in H27 cells. A ChIP-grade antibody against Histone3-acetylation (H3Ac), which commonly serves as a marker of open chromatin status, was used. With the treatment of Entinostat and Panobinostat, the enrichment for the euchromatin marker H3Ac were ∼1.7-fold and ∼2.9-fold higher than with no treatment group, respectively (**Figure 7**). Therefore, the HDAC inhibitors (Entinostat and Panobinostat) induced the euchromatin state in the CRISPR/Cas9 target region.

**Figure 7.**
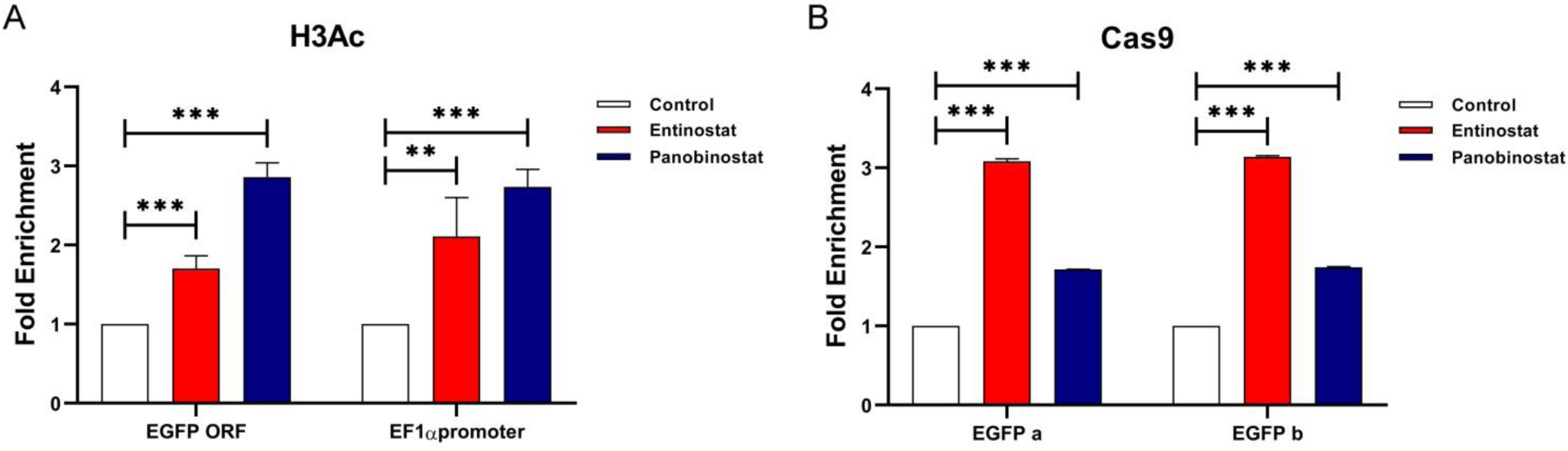
Chromatin immunoprecipitation (ChIP) and qPCR analysis of chromatin state and the binding between Cas9 and targeted DNA with HDAC inhibitors treatment. **A.** ChIP-qPCR was performed by using the antibody directly against open chromatin marks Histone3-acetylation (H3Ac). **B.** ChIP-qPCR was performed by using Cas9 antibody directly against the complex of Cas9 protein and the target loci of chromosome. The probed regions were located closely to CRISPR/Cas9 target sequences (−200bp, +200). Standard positive and negative controls have been described^10^.

To further determine the binding of Cas9 protein and target loci of chromosome in a direct manner, ChIP-qPCR assays were performed using a specific Cas9 antibody as well. The enrichment of Cas9 protein binding to target DNA was ∼3.0-fold and ∼1.7-fold higher than the group with no Entinostat or Panobinostat treatment (**Figure 7**). Of note, enrichment of Cas9 protein binding to targeted DNA was consistency with gene knock-in results (**Figure 6B**). Collectively, these results provide direct evidence for HDAC inhibitors improving the accessibility of chromatin and increasing the binding of Cas9 and targeted DNA (**Figure 8**).

**Figure 8.**
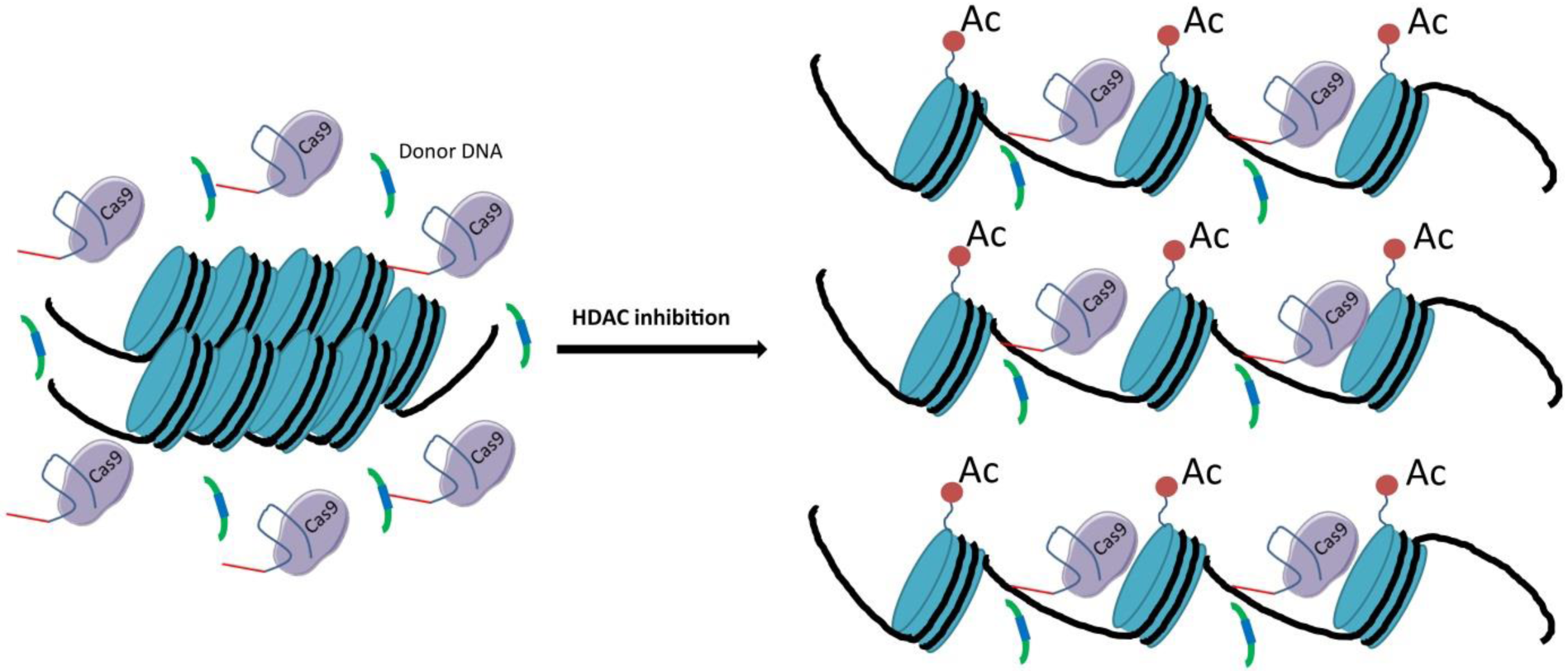
A proposed model of HDAC inhibition and CRISPR/Cas9 mediated gene editing. HDAC inhibition causes unwinding of the target DNA by the addition of acetyl groups to the histones, and thus improving the accessibility of the DNA for gene editing using the CRISPR/CAS9 system. This would enable binding of the nuclease to the desired target sequence and cut the DNA as well as HDR.

## Materials and methods

### Cell lines

HEK293T, HeLa and HT29 cells were purchased from American Type Culture Collection (ATCC, Wesel, Germany). H27^22^, HEK.EGFP^TetO.KRAB^ and HT29-EGFP is a single cell-derived clone from HeLa, HEK293 and HT29 for constitutively expressing EGFP, which have been described elsewhere^10,23,24^. All cells were cultured in DMEM medium (Gibco® by Life Technologies) supplemented with 10% Fetal Bovine Serum (FBS; Invitrogen, Breda, The Netherlands) and 1% Penicillin/Streptomycin (Gibco® by Life Technologies) at 37 °C with 5% CO2.

### Chemical reagents

The HDAC and HAT inhibitors Entinostat, RGFP966, C646, MG149, TMP195, PC134051 were purchased from Selleckchem. The purity of these inhibitors was assessed by Selleckchem (>99%). The Entinostat analogues have been described elsewhere^21^.

### Cell viability detection

An MTS assay was performed to determine the dose of the HDAC and HAT inhibitors that can be administrated to the cells without affecting cell viability. Cells (3×10^3^ per well) were cultured overnight and incubated with corresponding compounds in 96-well plates for 24 hr. The next day, cells were incubated at 37°C with MTS for 90 min following the instruction of CellTiter 96 AQueous One Solution Reagent (Promega, Madison, USA). The absorbance was measured at the wavelength of 490 nm by a Synergy H1 plate reader (BioTek, Winooski, USA). Experiments were performed in triplicate and repeated at least three times.

### Recombinant Plasmids

The construction of recombinant AdV shuttle plasmids has been detailed elsewhere^18^. Briefly, the recombinant plasmids contain phosphoglycerate kinase 1 gene promoter (PGK) and the simian virus 40 (SV40) polyadenylation signal. The pAdSh.PGK.Cas9 has a PGK and SV40 element for controlling the hCas9 expression. The Cas9 ORF was isolated from plasmid Addgene #41815. The gRNA expression units (gRNA-GFP) based on the U6 RNA Pol III promoter were retrieved from plasmids Addgene #41820. The constructs above were inserted into the MCS of pAdSh.MCS.SV40, resulting in the construct pAdSh.U6.gRNA^GFP^. The E1- and E2A-deleted fiber-modified AdV molecular clones pAdV^Δ2^P.Cas9.F^50^ and pAdV^Δ2^U6.gRNA^GFP^.F^50^ were assembled in BJ5183^pAdEasy-2.50^ *E. coli* via homologous recombination after transformation with MssI-treated AdV shuttle plasmids^25^.

For gene knock-in (EGFP-EBFP converting) experiments, the recombinant DNA has been detailed elsewhere^23^. Briefly, the gRNA expression plasmid AX03_pgEGFP and the hCas9 nuclease plasmid (Addgene plasmid 41815) were used for generating DSBs at the EGFP sequence. The AX63_pTHG was used as a donor template for converting EGFP into EBFP via HDR. The AM51_pgNT was served as a negative control.

### Plasmid transfection

The plasmids transient transfection using polyethyleneimine (PEI; Polysciences) has been detailed elsewhere^10^.

### pAdV^Δ2^P.Cas9.F^50^ and pAdV^Δ2^U6.gRNA^GFP^.F^50^ viral vector production, purification and titration

The production of viral vectors pAdV^Δ2^P.Cas9.F^50^ (AdV-Cas9) and pAdV^Δ2^U6.gRNA^GFP^((AdV-gRNA) have been described elsewhere^18,25,26^. Briefly, AdV particles were initiated by transfecting PER.E2A cells using PEI solution and PacI-linearized plasmids pAdV^Δ2^P.Cas9.F^50^ and pAdV^Δ2^U6.gRNA^GFP^.F^50^. After overnight incubation at 39°C, the culture media was replaced and cells were transferred to 34°C. The cells were harvested and following by three cycles of freezing and thawing in liquid nitrogen and in a 37°C water bath. Rescued AdVs presented in supernatants and amplified through propagation on PER.E2A cells newly seeded in a T75 flask. Large scale AdV produced in 16 T175 cell cultures flasks (Greiner Bio-One), following by CsCl gradient centrifugation method for purification. The titers of purified AdV were determined by TCID50 assay which has been detailed^18^.

### AdTL Viral vector production and purification

AdTL is a both E1- and E3-deleted recombinant serotype 5 adenovirus which contains a green fluorescent protein (GFP) and luciferase expression cassette^27^. THEK293 cells were seeded in 10 T175 flasks and transduced with AdTL in a dosage of 10 transduction units (TU)/cell to produce a large batch of viruses. When CPE of 100% was reached, the cells were harvested and centrifuged at 1000 rpm for 5 min. The pellet was subjected to three cycles of freezing and thawing using dry ice and a 37°C water bath, respectively. Subsequently, the suspension was centrifuged again at 4000rpm for 10 min. The supernatant was purified using a Q-sepharose-XL column for chromatography of adenovirus as described previously[29]. Separations were carried out using a flow rate of 2mL/min. The column was equilibrated using 25mL application buffer (50mM Tris-HCl pH 8,0 1.0mM MgCl2). Thereafter, the AdTL virus, dissolved in 5mL application buffer, was loaded on the column. The column was washed with 25mL application buffer followed by 25mL wash buffer (containing 0.3 M NaCl). The virus was eluted from the column using 5 mL elution buffer (containing 1M NaCl). The column was cleaned using 10mL 0.1 M NaOH followed by 25 mL demi H_2_O. An OD260 measurement was performed using the NanoDrop 1000 Spectrophotometer (ThermoFisher Scientific, USA) to determine the number of virus particles. In addition, a Limiting dilution assay with AdTL dilutions was performed to determine the TCID50. Furthermore, purification by the Q sepharose XL column was checked by loading non-purified and purified AdTL virus, mixed with SDS loading buffer (4x) on a 12,5% SDS-PAGE gel, which was run for 2 hr at 160 V. Subsequently, the gel was stained with Instant Blue for 30 min to make proteins visible. Furthermore, the purified virus was placed in a dialysis frame (Pierce) and dialyzed in dialysis buffer (10% glycerol, 10 mM Hepes, 1mM MgCl, pH=7.4) to remove the elution buffer, after which a final limiting dilution assay was performed.

### Transduction experiments

Cells were co-transduced with AdV.Cas9 (pAdV^Δ2^P.Cas9.F^50^) and AdV.gRNA-eGFP (pAdV^Δ2^U6.gRNA^GFP^) to measure whether HDAC/HAT inhibitors affect the quantity of EGFP gene knockout induced by CRISPR/Cas9. Cells were seeded with a density of 200,000 cells/well in a 6-well plate. After 24 hr, the medium was replaced by medium (5% FBS) containing HDAC/HAT inhibitors with indicated dose. After 24 hr, the cells were co-transduced with AdV.Cas9 and AdV.gRNA-EGFP in a quantity of 30 TU/well. All co-transductions were performed in a 1:1 ratio. Cells were subcultured for 12 days to remove EGFP protein from cells with disrupted EGFP ORFs. Subsequently, fluorescence microscopy, flow cytometry, western blot and the T7e1 assay were performed to assess which percentage of EGFP genes in cells had been cleaved by CRISPR/Cas9.

Furthermore, mock transduced H27 cells served as a control to determine the effect of the different compounds on the endogenous EGFP gene expression. 24 hr prior to treatment, H27 cells were seeded with a density of 200,000 cells/well. Subsequently, the cells were exposed to a medium containing the same concentrations of HDAC/HAT inhibitors as the knockout assay. After three days, flow cytometry was performed to determine the intensity of EGFP expression. HeLa and HT29 (without EGFP) cells were transduced with the AdTL virus, which encodes for GFP and luciferase, to determine the effect of the HDAC/HAT inhibitors on transgene expression levels. HeLa cells were seeded at a density of 400,000 cells/well in a 6-well plate. After 24 hr the medium was replaced by a medium containing indicated doses of HDAC/HAT inhibitors. After 24 hr incubation with the compounds, AdTL was added at a dosage of 30 TU/cell. Mock-transduced cells served as control. After three days, flow cytometry was performed on the AdTL transduced cells. When a difference in transgene expression was detected, an additional experiment was performed to clarify whether the effect was due to a difference in viral transduction or transcription of the transgene. This was determined by administration of the compound 24 hr prior to AdTL transduction or immediately after AdTL transduction.

### Gene-editing (NHEJ and HDR) based on EGFP-to-EGFP fluorochrome conversion using plasmid donor

HEK.TLR^TetO.KRAB^ cells were seeded at 2×10^5^ per well of 6-well plates overnight. Before DNA transfection, cells were treated with indicated compounds for 48 hr. The DNA transfection was performed by adding 1 mg/mL polyethyleneimine (PEI; Polysciences) with different plasmids including Cas9, pTHG. Donor and gRNA-EGFP-containing RGNs which have been detailed elsewhere^23^. The different transfection complex was replaced by regular culture medium in the presence or absence of Dox. The cells were sub-cultured circa every 3 days for up to 11 days. The frequencies of EGFP-negative and EBFP-positive cells cultured in medium with Dox or without Dox were determined by flow cytometry.

### HDAC1, 2 and 3 knockdown by siRNAs

For HDAC knockdown, H27 and HT29 cells were transfected using Lipofectamine 2000 (Invitrogen, Carlsbad, USA). Cells were seeded at 2×10^5^ per well of 6-well plates overnight. On the next day, cells were transfected with siRNAs (100 nM/well) against HDAC1 (MISSION, esiRNA HDAC1), HDAC2 (MISSION, esiRNA HDAC2) and HDAC3 (M-003496-02-0005, siGENOME). siRNAs for HDAC1 and HDAC2 were purchased from Millipore Sigma (Burlington, Massachusetts, USA). siRNAs for HDAC3 were purchased from GE Healthcare Dharmacon (Lafayette, Colorado, USA), or control siRNAs (Burlington, Massachusetts, USA). At least three gene-specific siRNAs were used for each gene silencing. After 3 days post-transfection, cells were lysed using RIPA buffer.

### RNA extraction and quantitative reverse transcriptase PCR (qRT-PCR)

Cells were washed with PBS and harvested by trypsin. RNA was extracted by the Maxwell LEV simply RNA Cells Kit (Promega, Madison, USA). RNA concentrations were determined by NanoDrop (Thermo Fisher Scientific, Waltham, USA). cDNA was synthesized using 200 ng total RNA by the Reverse Transcription kit (Promega, Madison, USA) according to the instruction. Primers are listed in Supplementary **Table S1**. Data analysis was processed by SDS v.2.3 software (Applied Biosystems, Foster City, USA).

### Fluorescence microscopy

Targeted EGFP knockout and AdTL transgene expression were monitored by fluorescence microscopy using a Zeiss Axiovert 25 CFL inverted light microscope (Carl Zeiss AG, Germany) with a 450-490 nm excitation and 515 nm emission filter set.

### Flow cytometric analysis

Cells were harvested and washed twice with standard FACS buffer (PBS plus 1% FBS). The proportion of GFP positive cells were quantified using FACS Calibur flow cytometer (BD, Franklin Lakes, USA). For EGFP and EBFP converting –mediated gene knock-in study, the BD FACSVerse cytometer (BD, Franklin Lakes, USA) was used. All the experiments were performed at the Central Flowcytometry Unit (University Medical Center Groningen). Data were analysed by FlowJo 7.2.2 software (Tree Star).

### Immunoblotting

Cells were washed with phosphate buffered saline (PBS) and harvested by trypsinization. Cells were lysed using RIPA buffer with Protease Inhibitor Cocktail (Thermo Fisher Scientific, USA). Protein concentrations were determined by a Pierce BCA Protein Assay Kit (Thermo Fisher Scientific, USA). Samples were separated by pre-cast SDS-PAGE (Bio-Rad, Hercules, USA) and transferred using a polyvinylidene difluoride membrane (PVDF). The PVDF membrane was blocked with 5% skimmed milk in PBST (0.1% Tween-20) at room temperature for 1 h and incubated at 4°C overnight with primary antibodies. Anti-rabbit or anti-mouse HRP conjugated antibodies were used to detect protein by chemiluminescence using ECL (Perkin Elmer, Western Lightning Plus ECL). Images were visualized by GeneSnap image analysis software (SynGene, Frederick, USA) and quantified with ImageJ software (National Institute of Health, USA). The following antibodies from Cell Signalling were used for immunoblotting, Cas9 (#14697), HDAC1 (#2062), HDAC2 (#5113) and HDAC3 (#2632). The dilution of primary antibodies was 1:1000 (v/v). The secondary antibodies rabbit anti-mouse (#P0260) and goat anti-rabbit (#P0448) HRP were purchased from Dako Denmark. The dilution of secondary antibodies was 1:2000 (v/v).

### Chromatin immunoprecipitation (ChIP) and qPCR

Chromatin immunoprecipitation (ChIP) and qPCR were performed to detect the alterations of chromatin state and the binding of Cas9 to targeted DNA in the presence or in the absence of HDAC inhibitors. The cells were cultured in the presence or absence of inhibitors at an indicated dose for 3 days, after which cell fixation was applied according to the protocol as described before^10^. Briefly, 2 ml of 11% formaldehyde solution were added to the cell culture medium. The culture flasks were agitated for 15 min at room temperature. Next, 1.1 ml of 2.5 M glycine was used to stop the fixation process. After a 5 min incubation at room temperature, cells were transferred to a 50ml tube. The harvested cells were subjected to centrifugation at 1350 ×g for 5 min at 4 °C for two cycles with PBS. Subsequently, the nuclei were isolated using pre-chilled solution (0.1%NP-40 in PBS) and collected by centrifuging at 10,000 rpm at 4 °C for 10 seconds. Next, to obtain DNA fragments, Micrococcal Nuclease (MNase) was used according to the manufacturer’s instruction. The ChIP-qPCR assays were carried out on 30 μg of cross-linked chromatin according to the HistonePath™ (Active Motif) ChIP-qPCR protocol. The ChIP-validated antibodies H3 pan-acetyl (Active Motif, cat # 39139) and Cas9 antibody (mAb) (8C1-F10) were used for the ChIP. Next, qPCR amplifications with primers targeting different regions (i.e. EF1α promoter, 5′ and 3′ EGFP gene segments) were performed. For Cas9 ChIP-qPCR, EGFP ORF primers were used. Additional primer pairs were used for the quality control of the ChIP-qPCR assays. All of the primer sequences have been detailed^10^.

The qPCR amplifications were performed in an ABI Prism 7900HT Sequence Detection System. The qPCR amplifications were performed in triplicate for each sample with the following procedures: a 2-min incubation period at 95°C, followed by 40 cycles with at 95°C for 15 seconds, at 58°C for 20 seconds at 72°C for 20 seconds. Next, the samples were incubated at 95°C, 55°C for 1 min, respectively. The binding events detecting every 1000 cells were calculated based on input amounts of chromatin, final ChIP volumes, and the primer efficiencies. Finally, the data were normalized according to the algorithm developed by Active Motif, Inc. (Carlsbad, CA), which is available as an online tool (https://www.activemotif.com).

### T7E1 assay

To determine the genome targeting efficiency of CRISPR/Cas9, T7 endonuclease 1 assay (T7e1) was performed. H27 Cells were harvested and genomic DNA was isolated using the (Qiagen, Germany) following the manufacturer’s instructions. The concentration of the isolated genomic DNA was determined using The NanoDrop One Spectrophotometer (ThermoFisher Scientific, USA). Thereafter, a PCR was performed using Taq polymerase (NEB, USA) with primers 5’ GAGCTGGACGGCGACGTAAACG 3’ and 5’ CGCTTCTCGTTGGGGTCTTTGCT 3’ for amplification (Sigma-Aldrich, Germany). The PCR amplification was as follows: an initial denaturation 95°C for 5 min, samples were subjected to 35 cycles of denaturation at 95°C for 30 seconds, annealing at 53°C for 30 seconds and elongation at 72°C for 40 seconds. Amplified DNA products were mixed with 1,5 μl NEB Buffer 2 and 3,0 μl nuclease-free water. An initial denaturation was performed following a ramp rate −2°C /second from 95°C and then −0.1°C /second from 85°C to 25°C. Subsequently, 1 μl T7e1 enzyme (NEB, USA) was added and incubated at 37°C in a water bath for 15 min. Gel electrophoresis was performed for detecting DNA fragments.

### Statistical analysis

The data were presented as mean ± SD (unless otherwise indicated). Data were derived from at least three independent experiments (unless otherwise indicated). Statistical analysis was performed by GraphPad software v.5.0 (La Jolla, CA, USA). Data were analyzed by a two-tailed unpaired student’s *t*-test (unless otherwise indicated). *p-values < 0.05; **p-values < 0.01; ***p-values < 0.001.

## Discussion

We comprehensively investigated the impact of HDACs and HATs on CRISPR/Cas9 mediated gene editing. We found that Entinostat (HDAC1/2/3 inhibitor) and Panobinostat (pan-HDAC inhibitor) enhanced Cas9 gene editing activity while other inhibitors (HDAC4/5/6/7/8/9) did not. We also found that RGFP966 (HDAC3 inhibitor) decreased CRISPR/Cas9 gene editing activity, which can be used for downregulation CRISPR/Cas9 gene editing activity. Importantly, Entinostat and Panobinostat dramatically increase gene knock-in rates. We confirmed these findings by knockdown of HDAC1, HDAC2 and HDAC3. We further identified that HDAC inhibition (Entinostat and Panobinostat) facilitated Cas9 access to target DNA and increased cutting frequencies. Our study revealed an essential role of HDACs in CRISPR/Cas9 mediated gene editing. We demonstrate that it is feasible to modulate CRISPR/Cas9 gene editing efficiency by regulating HDAC activity. This method might also be widely used in regulating other nucleases (ZFNs, TALENs, CRISPR/Cas9 and dCas9)-mediated genome and epigenome editing.

We showed that inhibition of HDAC1 or HDAC2, but not other HDACs or HATs, significantly enhances the gene knockout and knock-in frequencies by uncoiling the chromatin structure. Several recent studies showed that the chromatin structure and nucleosomes dramatically affect CRISPR/Cas9 mediated gene editing^10,28–30^. Therefore, we hypothesized that alteration of chromatin state by HDAC/HAT inhibitors can regulate genome editing by CRISPR/Cas9. Our data support this hypothesis and further show that HDAC1 and HDAC2 play an essential role in improving CRISPR/Cas9 mediated gene editing. Furthermore, we carried out an in-depth investigation to show that downregulation of HDAC1 or 2 activities can open the chromatin and improve the binding between Cas9 proteins and target DNA.

Our data revealed that HDAC inhibition shows remarkable enhancements of HDR events. The extremely low gene knock-in efficiency is one of the main obstacles for gene editing. Our findings contribute to the understanding gene editing by HDR in two aspects. Firstly, our study provides a new practical solution for enhancing gene knock-in with FDA proved HDAC inhibitors. Furthermore, our study suggests that histone acetylation and deacetylation may play an important role in HDR. One possible explanation is that the histone octamer complex around DNA may have strong steric effects for large DNA fragment insertion and integration by HDR. HDAC inhibitors cause unwinding of the target DNA by the addition of acetyl groups to the histones, and thus not only improves the accessibility of the targeted DNA to Cas9 protein, but also to the donor DNA for HDR. This enables binding of desired target sequence to the nuclease as well as to donor DNA. Thus, both knockout and knock-in can be enhanced. In addition, our results also indicate that the HDR events have a positive correlation to the binding of Cas9 protein and targeted DNA (Figure 6B and Figure 7B). Previous study has shown that dissociation of Cas9 protein from double-stranded DNA is ∼6 h which coincides with the HDR time ∼7 h^31,32^, we speculate that Cas9 protein may have interactions with double-stranded DNA and facilitate the HDR process during thisresidence time. Nonetheless, the precise mechanism of how Cas9 protein interacts with chromatinized DNA remains largely unknown. This needs to be clarified by future studies. Future studies may also need to clarify how Cas9 protein interacts with plain DNA (without histone) and nucleosomal DNA (with histone).

DSBs repair can also be affected by cell cycle arrest, which will eventually influence the gene editing process^19,20^. As HDAC inhibitors might affect cell cycle, we investigated the cell cycle changes with the treatment of HDAC inhibitors. We found that Panobinostat (pan-HDAC inhibitor) can increase the proportion of cells in G2/M phase, but we did not observe any cell cycle changes with Entinostat. Thus, we have excluded the gene editing process affected by cell cycle alterations induced by Entinostat. In addition, some of the HDAC or HAT inhibitors can regulate virus transgene expression. Our data has shown that Panobinostat significantly enhance Cas9 protein expression. Similarly, pan-HDAC inhibitors, such as Valproic Acid (VPA), can enhance ZFN expression and cell cycle modulation^33^. Also, another study has shown that pan-HDAC inhibition (trichostatin A and vorinostat) can be applied in transient IDLV-mediated ZFN expression modulation. However, we did not observe significant Cas9 expression changes in the condition of HDAC1, HDAC2 and HDAC3 specifically inhibition by Entinostat. Taken together, multiple factors, including G2/M phase cell cycle arrest and an increase in Cas9 expression, may contribute to CRISPR/Cas9 gene editing by Panobinostat. However, our study proved that gene editing efficiency enhanced by Entinostat is not due to those factors, but through changing the chromatin state.

Interestingly, inhibition of HDAC3 attenuated CRISPR/Cas9 gene editing activity, which suggests that HDAC3 played an extraordinary role in the gene editing process. It might be possible to use HDAC3 inhibitors for turning down the activity of Cas9. The first small molecular Cas9 inhibitors have been uncovered recently which can be used for inactivating Cas9 activity^34^. Although those inhibitors are with high potency, they might be only effective for SpCas9. However, HDAC3 inhibitors can be used for downregulating the gene editing efficiency through alteration chromatin state regardless of nucleases. HDAC and HAT inhibitors have shown mixed results in different studies and in the treatment of cancers^17^. Unlike HDAC1 and HDAC2, which only function in the cell nucleus, HDAC3 is able to shuttle between the nucleus and the cytoplasm^35,36^. Inhibition of HDAC3 in the cytoplasm or in the nucleus may have different effects on Cas9 mediated gene editing process. Furthermore, HDAC3 is present at DNA replication forks and inhibition of HDAC3 led to a significant reduction in DNA replication fork velocity^37^. Treatment of HDAC3 inhibitor caused inefficient or slowed DNA replication^37^. This inefficient DNA replication may affect the DSB repair induced by CRISPR/Cas9 and decrease the gene editing efficiency. Therefore, this unique feature of HDAC3 may be one of the explanations in the mechanism on the decreased gene editing efficiency by HDAC3 inhibition.

In conclusion, our study provides a practical option for improving gene editing efficiency through chromatin de-condensation using HDAC inhibition. Furthermore, our study facilitates a deeper understanding of gene editing process by altering the accessibility of the DNA through histone acetylation and histone deacetylation.

## Acknowledgements

We thank Manuel Gonçalves (Leiden University Medical Center, Dep. Cell and Chemical Biology) for providing us with the adenoviral vectors encoding Cas9 and a GFP-targeting gRNA, the human cell lines H27 and HEK.EGFP^TetO.KRAB^ and the plasmids AX03_pgEGFP, AM51_pgNT and AX63_pTHG.donor. We acknowledge Petra E. van der Wouden and Rita Setroikromo for technical support. We thank the Central Flowcytometry Unit of University Medical Center Groningen for providing technical support of flow cytometry. We thank Martijn Zwinderman for support of ChIP-qPCR study.

## Contributions

H.J.H. and B.L. designed the study. B.L. and S.W.C performed the HDAC screening, knockout and knock-in work and analyzed data. A.L.R. initially set up the AdV-CRISPR/Cas9 knockout system for HDAC screening with help of H.J.H. and B.L. D.C. performed Western blot, managed and analysed the figures with the help of B.L and S.W.C. qPCR and q-rtPCR were performed by S.W.C and D.C with the help of B.L. D.K. initially started the knock-in study with the help of B.L. S.W.C and M.A. tested Entinostat analogues with the help of B.L. D.J.F and F.Y.C provided Entinostat analogues and gave suggestions on HDAC inhibitors. B.L. wrote the manuscript with input from other authors. H.J.H supervised the project.

