## Supplementary Fig. S1-S4 and Table S1 for "Inhibition of Histone deacetylase 1 (HDAC1) and HDAC2 enhances CRISPR/Cas9 genome editing"

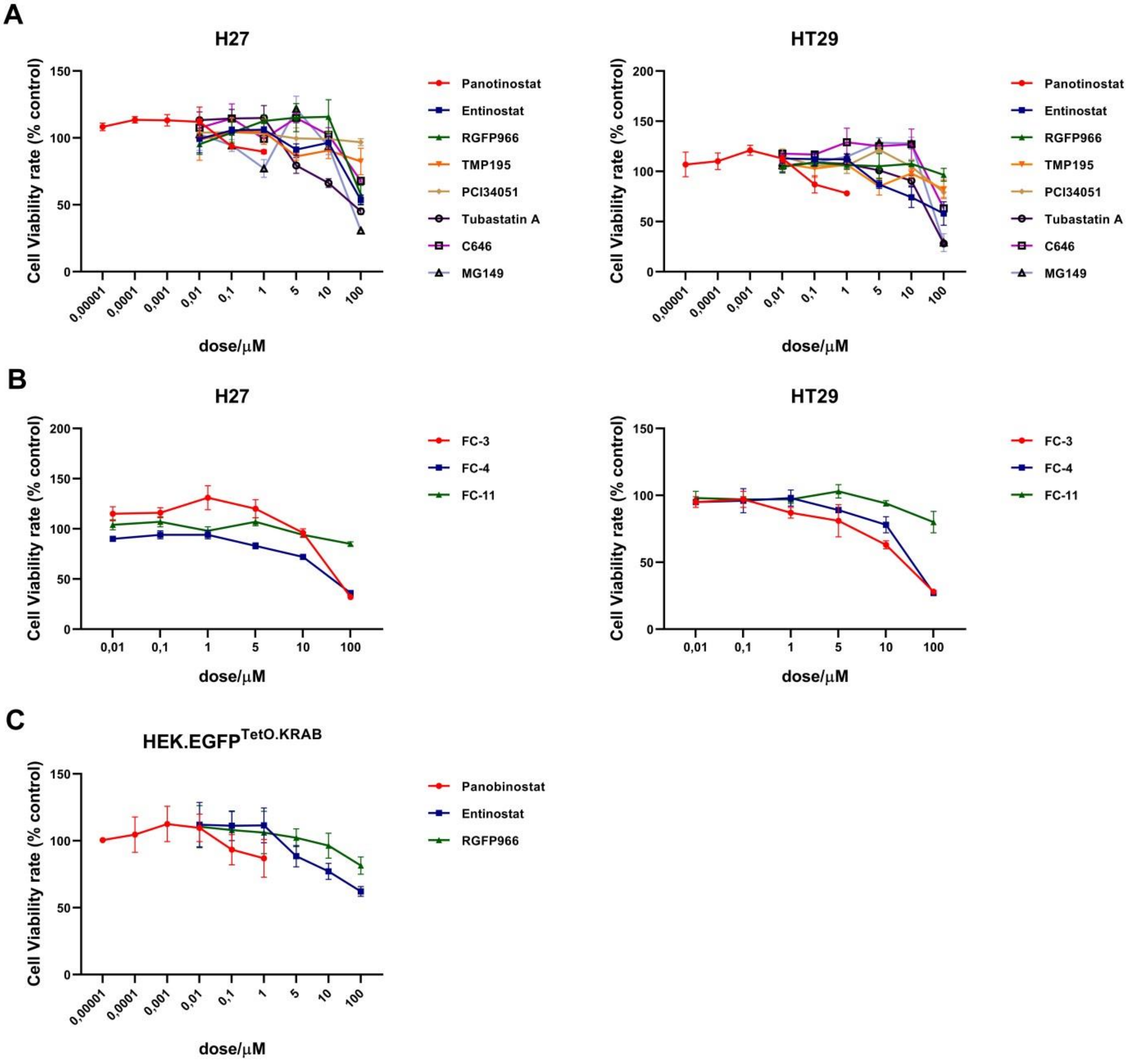


**Figure S1 Cell viability assessment with HDAC/HDAT inhibitors treatment.** Cells were treated with inhibitors at indicated dose for 24 h, and cell viability was determined by MTS assay. Data in bar graphs are represented as mean ± SD (n≥3).

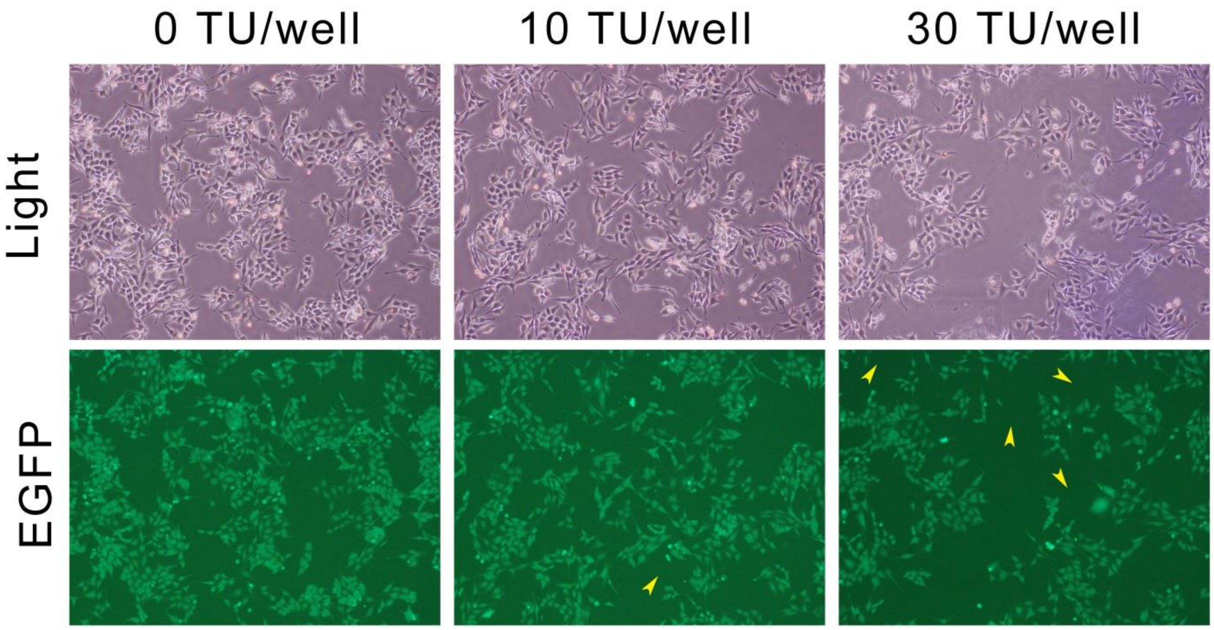


**Figure S2 A MOI of 30 TU/cells is suitable for AdV-Cas9 and AdV-gRNA co-transduction**. Fluorescence microscopy of H27 cells transduced with 0, 10, 30 or 100TU/well of both AdV-Cas9 and AdV-gRNA without fluorescence (upper panel) and with EGFP fluorescence (lower panel). The yellow arrows show the EGFP knockout cells.

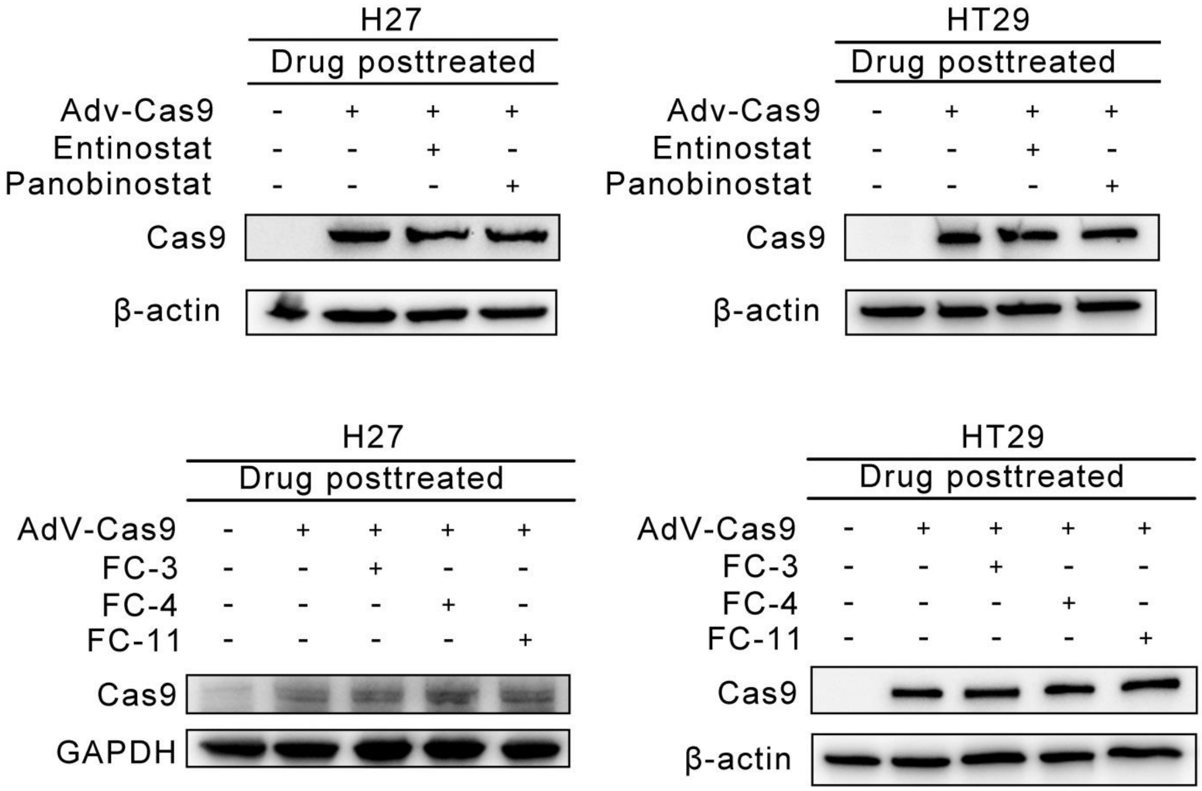


**Figure S3.** Western blot analysis of AdV-Cas9 protein expression with post-treatment of HDAC inhibitors using the same dose as shown in knockout for 48 hr.

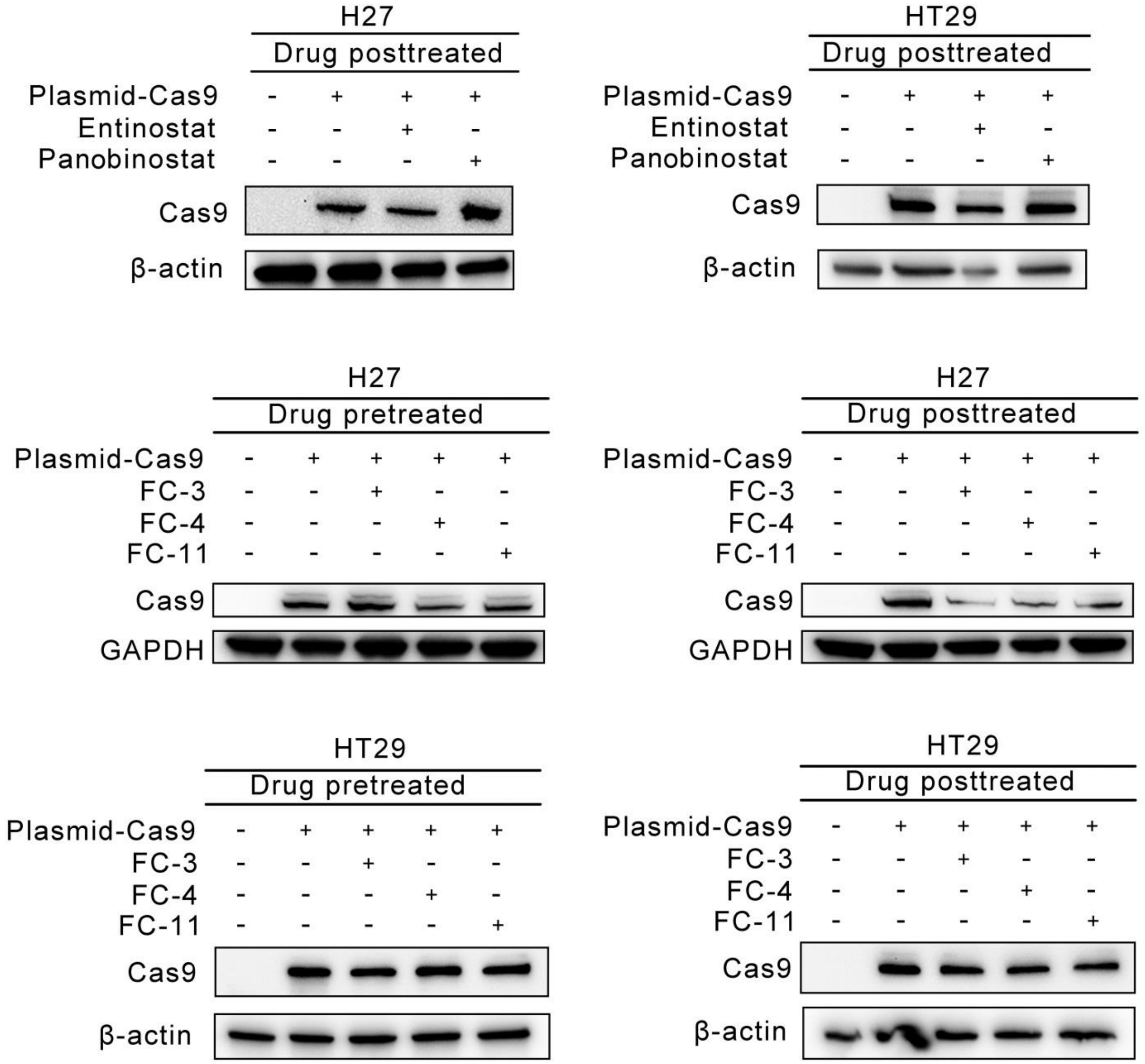


**Figure S4.** Western blot analysis of plasmid Cas9 protein expression with pre-treatment or post-treatment of HDAC inhibitors using the same dose as shown in gene editing study.

**Supplementary Table S1. List of primers used for HDACs RT-qPCR**

| Name | Strand | Sequence |
|---|---|---|
| HDAC-1 | F | 5’-GGAAATCTATCGCCCTCACA -3’ |
| | R | 5’-AACAGGCCATCGAATACTGG -3’ |
| HDAC-2 | F | 5’-AGACTGCAGTTGCCCTTGAT -3’ |
| | R | 5’-TGCGCAAATTTTCAAACAAA -3’ |
| HDAC-3 | F | 5’-TGGCTTCTGCTATGTCAACG -3’ |
| | R | 5’- CCCGGTCAGTGAGGTAGAAA-3’ |
| GAPDH | F | 5’-ACCCAGAAGACTGTGGATGG -3’ |
| | R | 5’-TCTAGACGGCAGGTCAGGTC -3’ |
